# A synapse-specific refractory period for plasticity at individual dendritic spines

**DOI:** 10.1101/2024.05.24.595787

**Authors:** Juan C. Flores, Karen Zito

## Abstract

How newly formed memories are preserved while brain plasticity is ongoing has been a source of debate. One idea is that synapses which experienced recent plasticity become resistant to further plasticity, a type of metaplasticity often referred to as saturation. Here, we probe the local dendritic mechanisms that limit plasticity at recently potentiated synapses. We show that recently potentiated individual synapses exhibit a synapse-specific refractory period for further potentiation. We further found that the refractory period is associated with reduced postsynaptic CaMKII signaling; however, stronger synaptic activation only partially restored the ability for further plasticity. Importantly, the refractory period is released after one hour, a timing that coincides with the enrichment of several postsynaptic proteins to pre-plasticity levels. Notably, increasing the level of the postsynaptic scaffolding protein, PSD95, but not of PSD93, overcomes the refractory period. Our results support a model in which potentiation at a single synapse is sufficient to initiate a synapse-specific refractory period that persists until key postsynaptic proteins regain their steady-state synaptic levels.

**Significance Statement:** A refractory period in which newly modified synaptic connections are unable to undergo further plasticity is a proposed mechanism through which newly formed memories can be preserved at the synaptic level while brain plasticity is ongoing. Here, we provide new insights into the spatiotemporal signaling mechanisms that regulate the establishment and maintenance of a refractory period for plasticity at individual excitatory synapses in the hippocampus, a region of the brain vital for learning and memory. Our results have implications in the identification of molecular targets that could serve to improve learning outcomes associated with disease.

## Introduction

Learning and memory are thought to rely upon long-term changes in neural circuit connections, both via alterations in the strength of existing synapses and through formation of new synapses^1–3^. One challenge for synaptic models of learning and memory has been how synapses can be both plastic, in order to encode new memories, and stable, in order to retain old memories^4–9^. A prominent hypothesis posits that synaptic connections that have recently been strengthened during learning are protected from further modification^10–12^. Indeed, there is evidence that when animals learn two distinct tasks over a close period of time, non-overlapping populations of synapses encode the two distinct tasks^2,13^.

An important component of synaptic learning models therefore would be mechanisms that support selection of non-overlapping sets of labile synapses to encode distinct tasks learned close in time. One such mechanism could be through the existence a period of time in which recently potentiated synapses are unable to undergo further potentiation; this type of metaplasticity has often been referred to as saturation of plasticity^4,10,14,15^. Indeed, in vitro studies at the circuit level have shown that recently potentiated hippocampal circuits are unable to undergo further potentiation within the first few hours after potentiation^16–19^. Furthermore, in vivo studies have shown that after inducing saturation of potentiation in the hippocampus of live rats, learning is impaired compared to control animals^14^. However, the cellular and molecular mechanisms that inhibit further plasticity at recently potentiated individual synapses, and the spatial and temporal scales over which they act, are not well understood.

Here, we show that prior potentiation at individual dendritic spines on dendrites of hippocampal CA1 neurons is sufficient to initiate a refractory period that prevents additional structural and functional plasticity. We further show that the refractory period is limited to synapses receiving prior potentiation, and that it is postsynaptically initiated and accompanied by reduced postsynaptic activation of CaMKII. Increasing the strength of the potentiating stimulus at 30 min is not sufficient to fully overcome the refractory period at individual synapses; however, it is fully released within 60 min. Finally, we show that increasing levels of the postsynaptic scaffolding protein PSD95, but not of PSD93, is sufficient to release the refractory period, such that previously potentiated synapses are again able to exhibit robust plasticity at a time when they would typically be refractory.

## Results

### Individual potentiated spine synapses are refractory to further plasticity

To test whether recently potentiated individual synapses on hippocampal CA1 neurons are restricted from further plasticity, our strategy was to induce long-term potentiation (LTP) at a single target spine and then test whether further potentiation could be induced at the same target spine 30 min later (**Fig. 1A**). After two baseline images, individual spines on EGFP-expressing CA1 neurons in slice cultures were stimulated with high-frequency uncaging of MNI-glutamate (HFU), which has been shown to induce concurrent long-term increases in synaptic strength and spine size^20,21^. As expected, following an initial HFU at time 0 (HFU_0_), we observed that stimulated target spines exhibited a long-term increase in size (**Fig. 1B, C, E**; 210% ± 20%; p < 0.001). Notably, an identical HFU stimulus at the same target spine 30 min later (HFU_30_) induced no further long-term growth (**Fig. 1B-E**; 102% ± 12%; p > 0.99). Importantly, previously unstimulated, size-matched (**Fig. S1A**) control spines on different dendrites of the same cells exhibited a robust long-term increase in size in response to HFU_30_ (**Fig. 1B, D, E**; 170% ± 18%; p < 0.01), supporting that the lack of plasticity observed in the re-stimulated target spine was not due to either (i) a decay in ability to undergo plasticity after 30 min in the bath chamber and/or (ii) the increased spine size following the first round of plasticity. Our results demonstrate that prior potentiation initiates a refractory period for further plasticity at individual dendritic spines.

**Figure 1.**
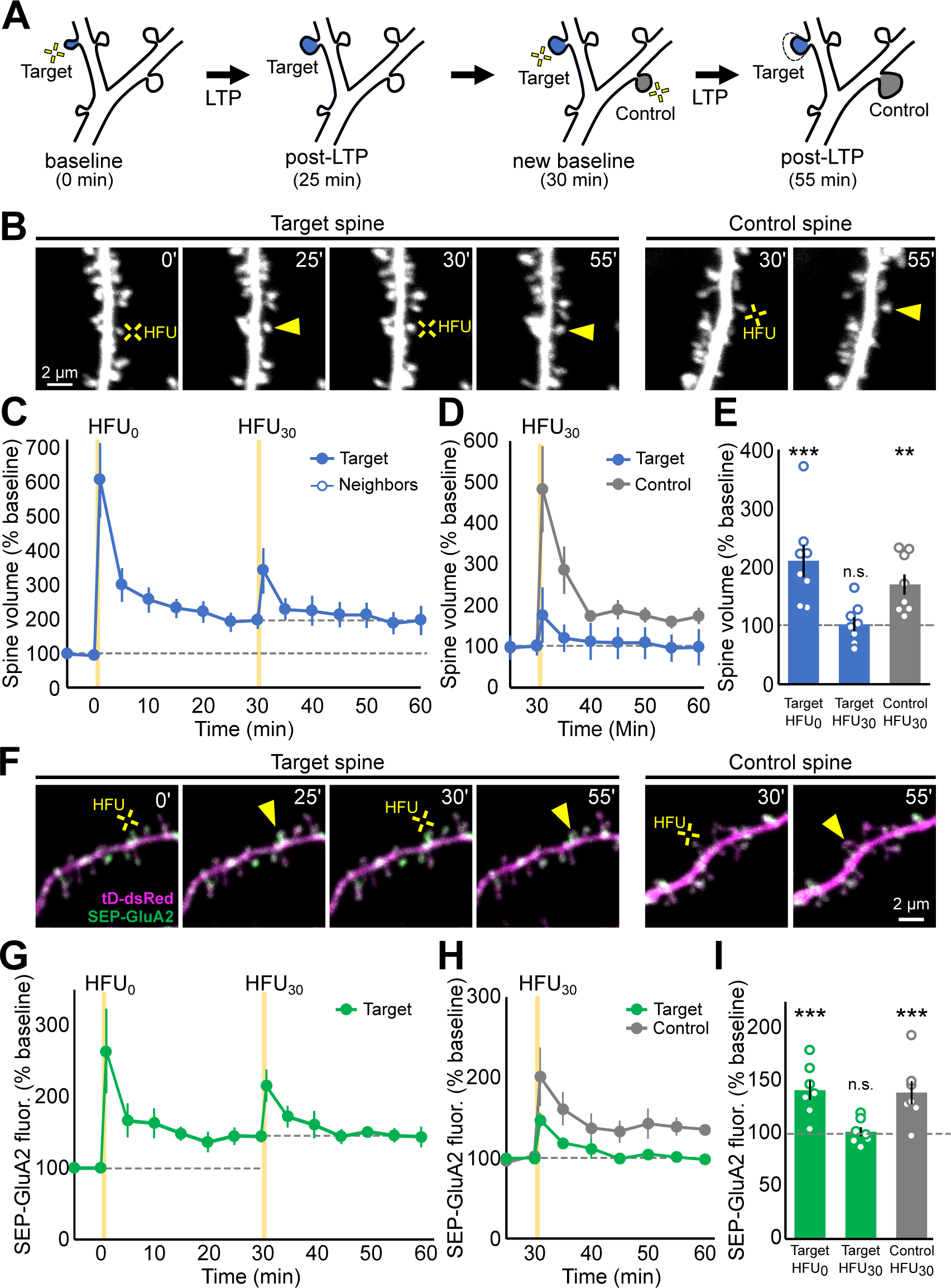
A single LTP-inducing stimulus initiates a synapse-specific refractory period for long-term growth and synaptic strengthening at individual dendritic spines. **(A)** Schematic of the experimental approach. An individual dendritic spine (target) was stimulated (yellow crosses) to induce LTP and then imaged every 5 min. At 30 min, the target spine was stimulated again along with a previously unstimulated, size-matched control spine (control) on the same cell. (**B**) Representative images of dendrites from an EGFP-transfected hippocampal CA1 pyramidal neuron directly prior to and 25 min after HFU_0_ and HFU_30_ at target (left images) and control (right images) spines. (**C-E**) The initial HFU stimulus (HFU_0_) resulted in long-term growth of target spines (filled blue circles/bars; n=8 spines/cells), but an identical HFU at 30 min (HFU_30_) did not drive additional growth. A size-matched control spine on the same cell grew in response to HFU_30_ (gray circles/bar; n=8 spines/cells). Neighboring spines were unchanged (open circles; n=73 spines/8 cells). Target spine data in **D** is from **C** re-normalized to the new baseline. (**F**) Images of dendrites from a CA1 pyramidal neuron transfected with SEP-GluA2 (green) and tDimer-dsRed (magenta) directly prior to and 25 min after HFU_0_ and HFU_30_ at target (left images) and control (right images) spines. (**G-I**) An initial HFU stimulus (HFU_0_) resulted in a long-term increase in surface expression of SEP-GluA2 in target spines (filled blue circles/bars; n=7 spines/cells), but an identical HFU at 30 min (HFU_30_) did not drive an additional increase. In contrast, HFU_30_ drove a long-term increase in surface SEP-GluA2 on a size-matched control spine on the same cell (gray circles/bar; n=7 spines/cells). Target spine data in **H** is from **G** re-normalized to the new baseline. Data represent mean ± SEM. Statistics: 2-way ANOVA with Bonferroni test. **p < 0.01, ***p < 0.001. See also **Figure S1**.

In addition to a long-term increase in spine size, potentiation at single spines also drives a long-term increase in surface AMPARs^20–22^. To examine whether insertion of surface AMPARs also exhibits a refractory period following potentiation at single dendritic spines, we transfected CA1 neurons with super ecliptic pHluorin (SEP)-tagged GluA2 to monitor surface expression of AMPARs^22–24^ along with tDimer-dsRed as a red cell fill (**Fig. 1F**). Importantly, by monitoring the red cell fill, we observed that the presence of SEP-GluA2 did not alter establishment of the refractory period for plasticity (**Fig. S1D-F**). By monitoring the SEP-GluA2 fluorescence, we observed that an initial HFU (HFU_0_) drove a long-term increase in surface GluA2 compared to baseline, as expected (**Fig. 1F, G, I**; 142 ± 9%, p < 0.001). An identical HFU stimulus at the same target spine 30 min later (HFU_30_) induced no further GluA2 insertion (**Fig. 1F-I**; 102 ± 4%, p > 0.99), despite that a second previously unstimulated, size-matched (**Fig. S1C**) control spine on a different dendrite of the same cell exhibited a robust long-term increase in GluA2 insertion in response to HFU_30_ (**Fig. 1F, H, I**; 139 ± 11%, p < 0.001). Thus, prior potentiation initiates a refractory period for both long-term spine growth and insertion of surface AMPARs.

### CaMKII activation is reduced in recently potentiated spines

To probe the molecular mechanisms underlying the refractory period following prior potentiation, we first focused on whether CaMKII signaling is altered in recently potentiated spines. CaMKII is a key integrator of synaptic activity and plays a key regulatory role in both LTP and LTD^25–28^. Using 2-photon fluorescence lifetime imaging (FLIM) of Camui-α, a genetically-encoded CaMKII activity reporter in response to a modified HFU protocol (see Methods) established in prior studies using this biosensor^29^, we imaged CaMKII activation (**Fig. 2A**) during and immediately following potentiation and saturation (**Fig. S2A-C**) of an individual dendritic spine. In response to the initial potentiating stimulation (HFU_0_), we saw a robust CaMKII activation, as detected by a sensor lifetime change of 120 ± 25 ps (**Fig. 2A, B, D**). Notably, a second identical potentiating stimulus 30 min later (HFU_30_) elicited a dramatically reduced change in CaMKII sensor lifetime (**Fig 2C, D**; 77 ± 14 ps, p = 0.029 compared to HFU_0_). Peak sensor lifetime change (**Fig 2C, D**; 130 ±14 ps) of size-matched control spines (**Fig. S2D**) was not different than the initial response (p = 0.65 compared to HFU_0_). Importantly, red fluorescence (**Fig. S2E**) and donor photon counts (**Fig. S2F**) in the target spines 25-30 min after HFU_0_ was not different from that of size-matched control spines, and initial spine size also did not influence the measured lifetime change during an HFU (**Fig S2G**). In sum, our results show that CaMKII activation is reduced in recently potentiated spines.

**Figure 2.**
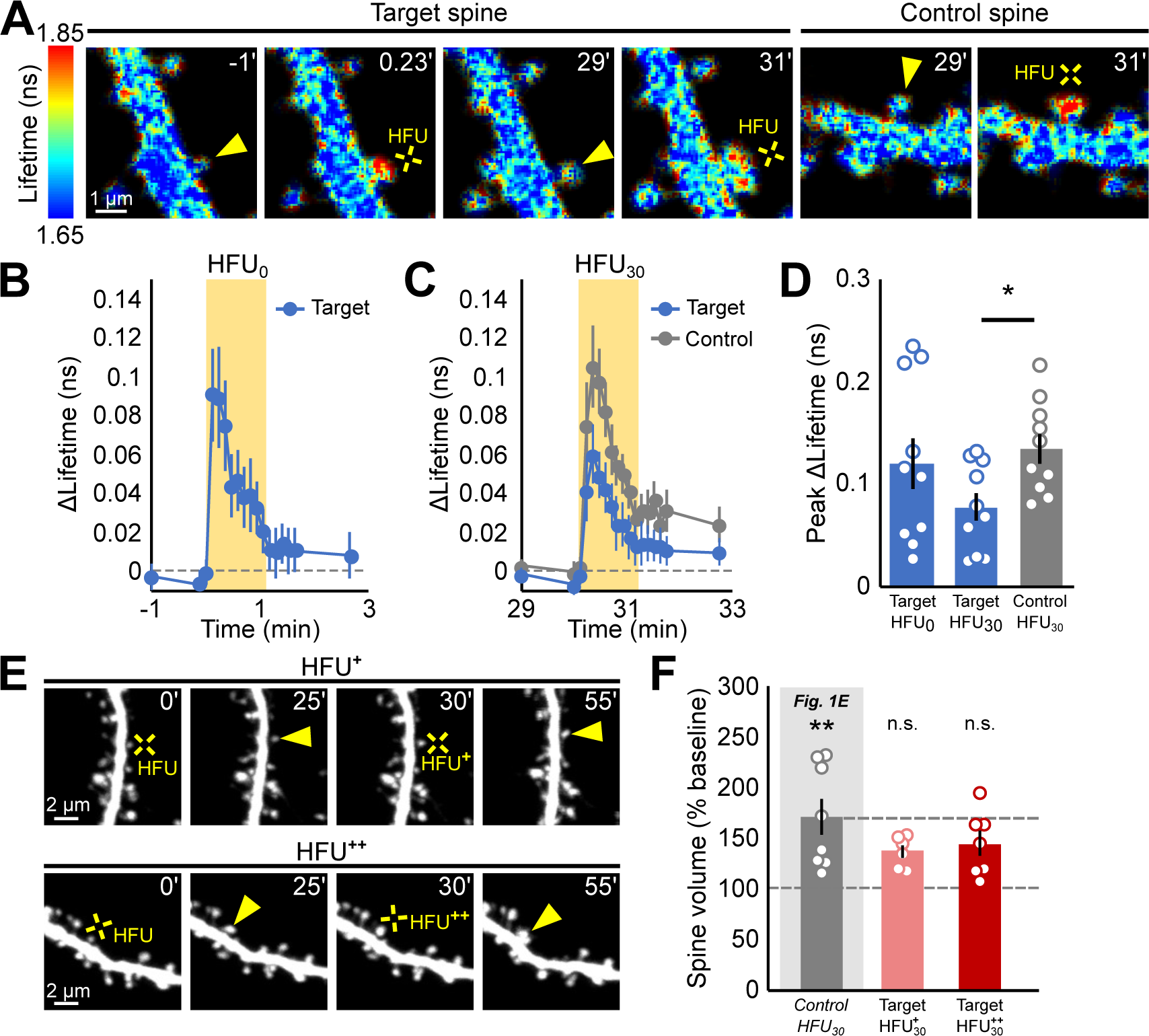
CaMKII activation is reduced in previously potentiated spines, but increased stimulation only partially restores plasticity. **(A)** Representative lifetime maps from neurons in slice culture expressing CaMKII FRET sensor prior to and during high frequency uncaging (HFU, yellow cross) at 0 min (HFU_0_) and 30 min later (HFU_30_) at target (left) and control spine (right). **(B)** The initial HFU stimulus (HFU_0_) resulted in robust activation of CaMKII at target spines (blue circles; n = 10 spines/cells). **(C, D)** CaMKII activation in response to the second HFU stimulus (HFU_30_) was reduced in previously potentiated target spines, but not in size-matched control spines (gray circles/bars; n = 10 spines/cells). Target spine data in **C** is from **B** re-normalized to the new baseline. **(E)** Representative images of spines before and after receiving HFU_0_ and a stronger HFU_30_^+^ (top) or HFU_30_^++^ (bottom). **(F)** Spines exhibited a non-significant trend toward long-term growth in response to HFU_30_^+^ (pink bars; n = 6 cells/spines; p = 0.18) and HFU_30_^++^ (red bars; n = 7 cells/spines; p = 0.06). Control HFU_30_ data are replotted from Figure 1E. Data represent mean ± SEM. Statistics: paired Student’s t-test used in **D**; 2-way ANOVA with Bonferroni test used in **E**. *p <0.05, **p < 0.01. See also **Figure S2**.

### Refractory period for plasticity at previously potentiated spines is only partially released increasing stimulus strength

Because we found that CaMKII activation was decreased in recently potentiated spines, we tested whether plasticity could be recovered by increasing the strength of the second stimulus to drive stronger activation of CaMKII. We therefore increased the strength of the HFU_30_ stimulus by increasing the duration of glutamate uncaging pulses from our standard 2 ms to a longer 4 ms (HFU_30_^+^) or 5-6 ms (HFU_30_^++^). In response to the stronger 4 ms HFU_30_^+^ stimulation at 30 min, target spines showed a trend toward growth (**Fig 2E, F**; 136 ± 6% p = 0.17). Further increasing the strength of the second stimulation to the even stronger 5-6 ms HFU_30_^++^ stimulation resulted in a slightly further increased, but non-significant long-term growth (**Fig 2E, F**; 144 ± 12%, p = 0.06). Further increasing the stimulus was not possible without damaging cell health (data not shown). Notably, despite the increase in stimulus strength with HFU_30_^+^ and HFU_30_^++^, the size increase of the target spine did not reach the magnitude or robustness that would be expected from a previously unstimulated control spine receiving standard stimulation (**Fig. 1E**, reproduced in **Fig. 2F**). Our results show that increasing the strength of second potentiating stimulus can partially overcome the refractory period; however, it does not fully recover plasticity to the level of spines which had not experienced prior stimulation.

### Refractory period for single spine plasticity is released within 60 min

Because a stronger stimulus was unable to fully recover plasticity of spines during the refractory period, we wondered if full recovery from the refractory period relied on a time-dependent restoration of a vital signaling process at individual spine synapses. Prior studies at the circuit level have shown recovery of saturation of plasticity within 1-2 hrs^16,18,19^, but some have attributed recovery to time-dependent alterations is synaptic ultrastructure^30^, while others have suggested that recovery does not occur at individual synapses, but instead can be attributed to the addition of new synapses^18^. We therefore examined whether recovery of plasticity could be observed at individual hippocampal CA1 synapses with a longer time interval.

To test whether increasing the time interval between the first HFU and the second HFU would permit recovery of plasticity, we repeated our experiments now with a 60 min interval between the two HFU stimuli (**Fig. 3**). As expected, target spines exhibited a long-term increase in size in response to the HFU_0_ stimulus (**Fig. 3A, B, F**; 171% ± 13%, p < 0.001). In contrast to the lack of additional plasticity observed at 30 min, a second identical HFU stimulus at 60 min (HFU_60_) resulted in an additional long-term increase in size of target spines (**Fig. 3B, D, F**; 135% ± 11%, p < 0.01). Importantly, recovery of plasticity was not due to secondary effects caused by a decay in size of target spines back toward their initial value over the 60 min interval, as the long-term increase in size induced by HFU_0_ was maintained for the full 60 min prior to HFU_60_ (20-30 min: 171 % ± 13%; 55-60 min: 166% ± 17%; p = 0.8). Furthermore, previously unstimulated, size-matched (**Fig. S3A**) control spines on a different dendrite of the same cells grew to a comparable magnitude in response to HFU_60_ (**Fig 3A, D, F**; 137% ± 9%, p < 0.01), supporting complete recovery of plasticity in the target spine at 60 min.

**Figure 3.**
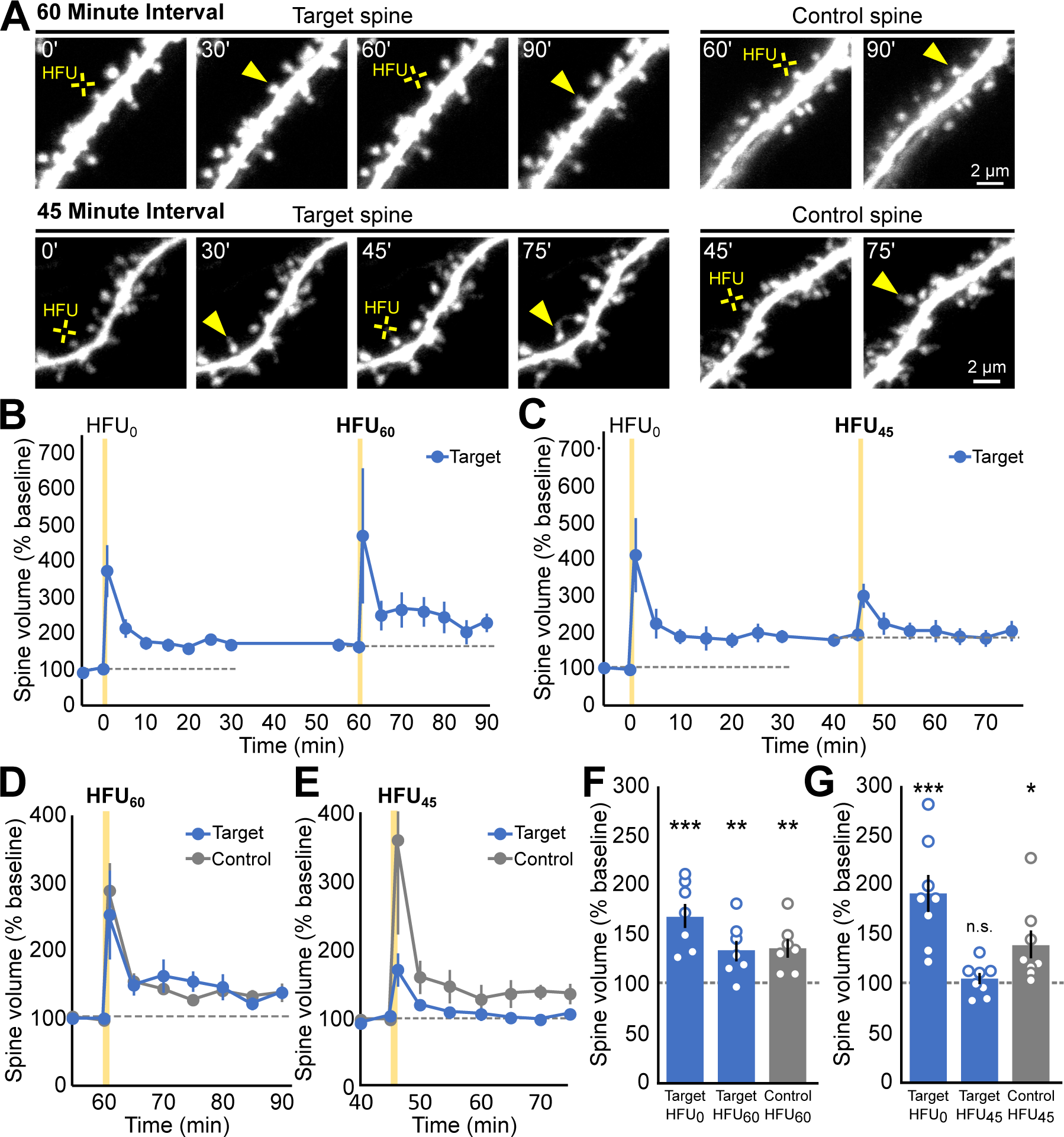
Refractory period for plasticity at previously potentiated spines is released within 60 min. **(A)** Images of dendrites from an EGFP-transfected CA1 neuron in slice culture directly prior to and 30 min after HFU_0_ and HFU_60_ at target and control spines (top images) or after HFU_0_ at and HFU_45_ at target and control spines (bottom images). **(B,D,F)** A second HFU at 60 min (HFU_60_) resulted in additional growth (filled blue circles/bars, n=7 spines/cells), comparable to that observed from size-matched control spines on the same cell (gray circles/bar; n=7 spines/cells). Target spine data in **D** is from **B** re-normalized to the new baseline. **(C,E,G)** A second HFU at 45 min (HFU_45_) did not result in additional growth of target spines (blue circles/bars; n = 8 cells/spines), despite that size-matched control spine on the same cell grew (gray circles/bars; n = 7 cells/spines). Target spine data in **E** is from **C** re-normalized to the new baseline. Data represent mean ± SEM. Statistics: 2-way ANOVA with Bonferroni test. *p <0.05, **p < 0.01. See also **Figure S4**.

To more precisely define the recovery time interval, we repeated our experiments with a 45 min interval between the two HFU stimuli. As expected, we observed that spines grew in response to HFU_0_ (**Fig. 3C, G**; 189% ± 19%; p < 0.001). However, with this interval, the target spine did not grow in response to HFU_45_ (**Fig. 3C, E, G**; 103% ± 6%; p = 0.80). In contrast, the size-matched control spine (**Fig. S3B**) grew in response to HFU_45_ (**Fig. 3E, G**; 138% ± 14%; p = 0.01). Our results support that the target spine was completely released from the refractory period for further plasticity between 45-60 min.

### Increased PSD95 expression level is sufficient to release the refractory period for plasticity at previously potentiated spines

Our data support the existence of a refractory period for plasticity at single spines that is released between 45 and 60 min after potentiation. In addition, because our glutamate uncaging stimulus bypasses presynaptic vesicle release, our results support that the refractory period is initiated via postsynaptic mechanisms. Ultrastructural studies have demonstrated that the PSD of potentiated spines does not increase in size for the first 30 min after LTP induction, but does so within 2 hrs^31^, suggesting that delayed PSD enlargement at potentiated synapses could contribute to limiting plasticity^30^. Furthermore, live imaging studies have shown that the synaptic expression level of several GFP-tagged postsynaptic scaffolding proteins does not increase within 30 min of LTP induction, despite a rapid increase in spine volume^32^. We therefore hypothesized that acceleration of the post-LTP PSD expansion through increasing the availability of key PSD proteins might allow for faster recovery from the refractory period. To test our hypothesis, we manipulated the expression level of PSD95, one of the most abundant PSD scaffolding proteins^33,34^ with important roles in regulating spine stabilization^35–39^, and which increases at newly potentiated spines only after a delay^40^.

To determine whether increased expression of PSD95 would accelerate recovery from the refractory period, we selected a 45 min time interval between the two plasticity-inducing HFU stimuli - a time at which the refractory period persisted, but such that synaptic molecular configuration would be close to recovered, and therefore increasing the expression of a single PSD protein might expedite the recovery of plasticity. CA1 neurons were transfected with GFP-tagged PSD95α^41^ (PSD95-GFP) and a red cell fill (**Fig. 4A**, top row). As expected, target spines exhibited a long-term increase in size in response to HFU_0_ (**Fig. 4B, C**; 180 ± 19%, p < 0.001). Notably, target spines with excess PSD95 exhibited an additional long-term increase in size in response to the second HFU at 45 min (**Fig. 4B, C**; 136 ± 10%, p < 0.05) that was comparable to that of previously unstimulated, size-matched (**Fig. S4A**) control spines (**Fig 4A, C**; 140% ± 13%, p < 0.05), suggesting complete recovery of plasticity in the target spine. Importantly, the magnitude of long-term spine growth in response to HFU_0_ (**Fig. S4C**) was not altered in cells with excess PSD95. Furthermore, increased synaptic AMPAR currents associated with excess PSD95^42,43^ did not contribute to overcoming the refractory period, because even in the presence of NBQX to block AMPAR currents, target spines exhibited long-term growth in response to HFU_45_ (**Fig. 4D**; 148 ± 14%, p < 0.05), comparable to that of previously unstimulated, size-matched (**Fig. S4B**) control spines. Thus, increased expression of PSD95 is sufficient to shorten the refractory period for plasticity induced by prior potentiation at single spines.

**Figure 4.**
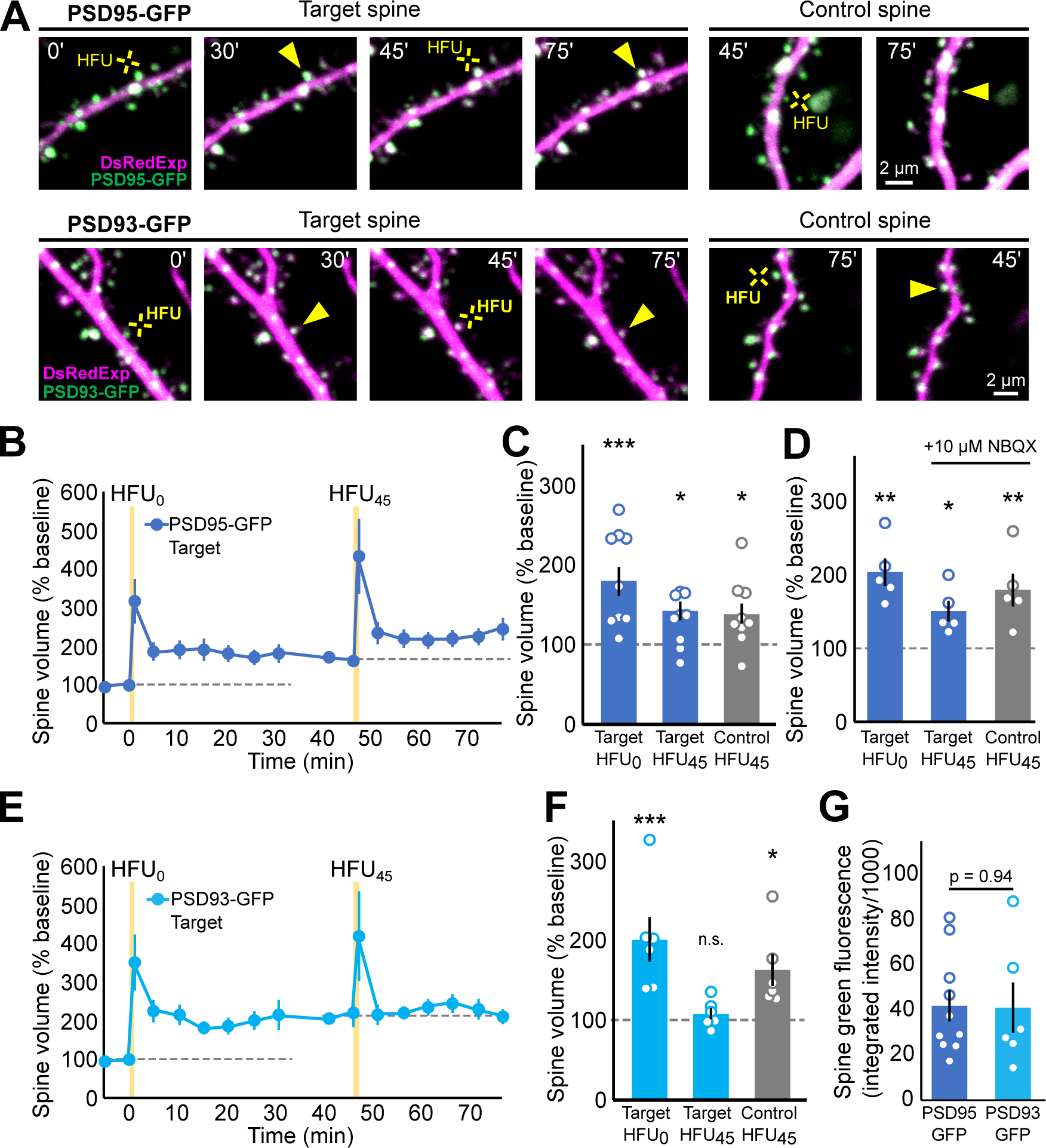
Increased availability of PSD95, but not PSD93, restores plasticity to spines in the refractory period. **(A)** Images of dendrites from CA1 pyramidal neurons transfected with DsRedExpress (magenta) and PSD95-GFP (green; top row) or PSD93-GFP (green; bottom row) directly prior to and 30 min after HFU_0_ and HFU_45_ at target and control spines. **(B,C)** PSD95-GFP-expressing target spines grew in response to a second HFU at 45 min (HFU_45_) (dark blue circles/bars; n=10 spines/cells, p < 0.05), even when (**D**) NBQX was added to block current through AMPARs (n = 5 spines/cells). **(E, F)** In contrast, PSD93-expressing target spines did not grow in response to HFU_45_ (light blue circles/bar; n=6 spines/cells; p > 0.99). **(G)** Expression levels of PSD95-GFP and PSD93-GFP were comparable. Data represent mean ± SEM. Statistics: 2-way ANOVA with Bonferroni test used in **C**, **D**, **F**; unpaired Student’s t-test used in G. *p <0.05, ***p < 0.001. See also **Figure S4**.

### Increased PSD93 expression level does not release the refractory period for plasticity at previously potentiated spines

Because PSD95 and PSD93 have extensive sequence identity (∼70% overall^44^), especially in the 3 PDZ binding motifs (86%), but have distinct functional roles^45,46^, we asked whether increased PSD93 levels would also shorten the refractory period for plasticity at single spines. We therefore repeated the experiment with GFP-tagged PSD93α^42^ (PSD93-GFP; **Fig. 4A**, bottom row). In contrast to the rescue of plasticity observed with PSD95-GFP, spines from cells with excess PSD93-GFP failed to exhibit long-term growth in response to a second HFU stimulus at 45 min (**Fig. 4E, F**; 109 ± 7%; p > 0.99), despite that previously unstimulated, size-matched (**Fig. S4D**) control spines exhibited robust growth (**Fig. 4F**; 164 ± 20%, p < 0.05). Importantly, spine expression of GFP-tagged proteins and initial size of the target spines were not different between PSD95- and PSD93-expressing spines (**Fig. 4G, S4E-G**). Altogether, our data show that increased expression of PSD95, but not PSD93, overcomes the refractory period for plasticity at single spines.

## Discussion

That synapses which have experienced recent plasticity become temporarily resistant to further plasticity has been proposed as a mechanism through which newly formed memories can be preserved at the synaptic level while brain plasticity is ongoing^4–8,10^. Here, we provide insights into the cellular and molecular mechanisms that regulate the establishment and duration of a refractory period for plasticity on dendrites of hippocampal CA1 neurons. We show that potentiation at single synapses is sufficient to establish a refractory period for further potentiation that is synapse-specific, lasts between 45-60 minutes, is initiated postsynaptically and accompanied by reduced postsynaptic CaMKII signaling, and is regulated by the expression level of the postsynaptic scaffolding protein, PSD95, but not PSD93.

### Initiation, time-course, and synapse-specificity of refractory period for potentiation

We demonstrate that a single LTP-inducing glutamatergic stimulus at an individual spine is sufficient to initiate a refractory period for further structural and functional potentiation. Earlier studies established that this type of metaplasticity, or ‘saturation of plasticity’ is observed at the circuit level^16–19^. Here, we show that the signaling mechanisms needed to establish the refractory period can be activated locally at single synapses, and do not require activating multiple axons as in circuit level studies using a variety of protocols to induce LTP, including tetanic, theta-burst, and pairing stimulation^16–19^. Importantly, because our glutamate uncaging stimulation bypasses presynaptic vesicle release, our results point to a postsynaptic locus of initiation for the refractory period. We further show that the refractory period for further potentiation is restricted to the potentiated spine; it is not observed at previously unstimulated, size-matched control spines on a different dendrite of the same cell, suggesting that the refractory period is synapse-specific. Although our studies do not exclude that immediate neighbors of potentiated spines also exhibit a refractory period for potentiation, this is unlikely as other studies have shown that LTP at an individual target spine lowers, rather than increases, the threshold for plasticity at spines that are within 10 µm of the potentiated spine^47^.

Because our experimental paradigm focused on single synapses, we were able to distinguish that full recovery can occur at these hippocampal CA1 synapses without the need to recruit additional naïve synapses^18^. It is likely that the success of induction and duration of the refractory period will be dependent on the pattern, strength, and extent of synaptic activity at single synapses. Indeed, it has been recently reported that different naturalistic patterns of activity influence the duration of long-term synaptic structural plasticity^48^.

### Molecular mechanisms of refractory period for potentiation

We show that CaMKII activation is reduced in recently potentiated spines, relative both to initial activation of the same spines following induction of LTP, and also to size-matched control spines on different dendrites of the same neuron. How might CaMKII activation levels be reduced specifically in recently potentiated spines? It is possible that the reduction in CaMKII activity is due to the local upregulation of endogenous CaMKII inhibitors, such as CaMKII inhibitor 1 (CaMK2N1), which has been shown to be upregulated after the induction of LTP^49^. Another possibility would be through mechanisms that drive reduced calcium influx following LTP. For example, a local feedback loop between the NMDA receptor and the Ca^2+^-activated small conductance potassium channel, SK2, could lead to a local reduction in the NMDAR Ca^2+^ currents^50^. Alternatively, activity^51–53^ and the induction of LTP^54^ have been shown to drive a rapid change from GluN2B-to GluN2A-containing NMDARs, which carry less Ca^2+^ current and are less favorable to induction of LTP^54,55^. This switch is likely to occur also at single synapses, as prolonged inactivation of synaptic transmission at individual synapses on cultured hippocampal neurons induces the opposite switch from GluN2A to GluN2B^56^. However, in at least some reports^56^, the LTP-induced subunit switch shows no recovery within one hour and it is not observed in slices from older animals comparable in age to those used for our experiments.

We found that increased levels of the postsynaptic scaffolding protein PSD95 restored plasticity to recently potentiated synapses in the refractory period. Our results suggest that low levels of PSD95 in recently potentiated synapses contribute to establishment or maintenance of the refractory period. Indeed, ultrastructural studies following the induction of LTP have demonstrated that the PSD takes between 30 min and 2 hrs to grow^31^, and molecular imaging studies have established that PSD-scaffolding molecules, including PSD95, accumulate to steady-state levels only after a delay of more than 30 min^32^. Intriguingly, we show that increased expression of PSD93, which is also at low levels following LTP, is not sufficient to restore plasticity to recently potentiated synapses. This might be viewed as surprising, as these two PSD-MAGUKs have a high degree of sequence identity, and share 3 PDZ domains, an SH3 and a GK domain with 2 palmitoylation sites^57^, and both have demonstrated roles in spine stabilization^35–39^. However, despite the similarities, PSD95 and PSD93 have many distinct physiological roles. For example, homeostatic upscaling of synaptic currents requires both PSD95 and PSD93, while scaling down does not require PSD93^45^. In addition, loss of PSD95 prevents maturation of silent synapses, while loss of PSD93 has the opposite effect, instead leading to accelerated maturation of silent synapses^46^. Furthermore, in nascent dendritic spines, PSD93 is enriched to mature levels within several hours, whereas PSD95 takes over 12 hours to reach mature levels, suggesting sequential roles in nascent spine stabilization^58^.

How might PSD95 levels regulate the refractory period? Our results provide experimental evidence in support of models based on ultrastructural studies positing that delayed expansion of the PSD limits further potentiation at recently potentiated synapses^30^. In these models, it has been proposed that the refractory period ends via the expansion of nascent zones, regions of the presynaptic bouton that lack glutamate release machinery but are apposed to the PSD^30^. In these models, synaptic strengthening requires that presynaptic nascent zones are converted into active zones by the addition of presynaptic vesicle release machinery, and only with time do nascent zones reappear through expansion of the PSD, again allowing plasticity at these synapses. Our results are consistent with a model in which excess PSD95 is sufficient to drive expansion of the PSD and addition of nascent zones, and thus recovery of plasticity. Molecularly, reduced PSD95 levels following LTP would be expected to permit increased levels of STEP_61_, a tyrosine phosphatase that targets GluN2B for endocytosis^59^, and thus would act to reduce the efficacy of LTP until the return of PSD95, exclusion of STEP_61,_ and subsequent return of GluN2B levels^51^.

### Implications for learning of refractory period for plasticity

A refractory period for plasticity during learning at recently potentiated synapses would serve an important role to exclude those synapses from incorporation into distinct subsequently-learned tasks, thus prevent overwriting of recently established memories still in the labile phase^4–8,10^. Another consequence of a refractory period on learning would be to temporarily prevent synapses from undergoing further potentiation related to the same task. Indeed, there is considerable evidence that repeated learning bouts activate the same sets of synapses^2,60,61^. Several have proposed that a refractory period for plasticity could be the basis for increased success of a type of learning known as spaced learning, in which breaks are incorporated into learning sessions^18,30^. Such breaks during spaced learning could serve to allow synapses to recover their ability to undergo plasticity. Indeed, spaced learning approaches have been shown to improve learning outcomes over traditional learning in which repetitions are temporally clustered in both humans and rodents^60,62,63^, and to improve learning under conditions where learning is challenged in disease^64,65^. It follows that manipulation of the molecular signaling mechanisms that regulate the refractory period could serve to improve learning outcomes associated with disease.

## Materials and Methods

### Preparation and transfection of organotypic hippocampal slice cultures

Cultured hippocampal slices (300-400 µm) were prepared from P6-P8 C57BL/6 mice of both sexes, as described^66^, and as approved by the UC Davis Institutional Animal Care and Use Committee. Slices were transfected at 9-15 DIV using biolistic gene transfer (180-210 psi), as described^67^. 6-8 mg of 1.6 µm gold beads (BioRad) were coated with 15 µg of EGFP-N1 (Clontech), or 10 µg of tDimer-dsRed together with 16 µg SEP-GluA2^24^, or 10 µg of DsRedExpress alone, or 10 µg DsRedExpress (Clontech) together with 1-2 µg of GFP-tagged PSD95α^41^ or PSD93α^42^. Slices were transfected 2-3 days (EGFP), 3-4 days (SEP-GluA2/tDimer-dsRed), or 24 hrs (PSD95/93/DsRedExpress) prior to imaging.

### Time-lapse two-photon imaging

Transfected CA1 pyramidal neurons at depths of 10–50 µm in slice cultures (11-17 DIV) were imaged using a custom two-photon microscope^68^ controlled with ScanImage^69^. Image stacks (512 x 512 pixels; 0.02 µm/pixel) with 1 µm z-steps were collected. For each neuron, one segment of secondary or tertiary basal dendrite was selected under epifluorescence and imaged at 5 min intervals at 29°C in recirculating ACSF (in mM): 127 NaCl, 25 NaHCO_3_, 1.2NaH_2_PO_4_, 2.5 KCl, 25 D-glucose, aerated with 95% O_2_/5% CO_2_, 310 mOsm, pH 7.2, with 0.001 TTX, 0 Mg^2+^, and 2 Ca^2+^. MNI-glutamate (2.5 mM; Tocris) was added at least 15 min prior to uncaging stimulation.

### High frequency uncaging (HFU) of glutamate

For all experiments except FLIM: HFU consisted of 60 pulses (720 nm; ∼7.5–9.5 mW at the sample) of 2 ms duration (4 ms for HFU^+^, 5-6 ms for HFU^++^) at 2 Hz delivered in ACSF containing (in mM): 2 Ca^2+^, 0 Mg^2+^, 2.5 MNI-glutamate, and 0.001 TTX. For FLIM: HFU consisted of 30 pulses (720 nm; ∼4 mW at the sample) of 4-6 ms duration at 0.5 Hz delivered in ACSF containing (in mM): 4 Ca^2+^, 0 Mg^2+^, 4 MNI-glutamate, and 0.001 TTX. Because earlier studies established that spine size can influence the magnitude of potentiation^20^, we selected target spines with an initial size range in the 25-50% quartile of sizes. The laser beam was parked at a point ∼0.5–1 µm from the spine head in the direction away from the dendrite. Cells with no noticeable transients in response to HFU_0_ were discarded and not further pursued.

### Image analysis

Images were analyzed using a custom MATLAB software, as described^68^. In brief, following application of a 3 x 3 median filter, background-subtracted integrated green and/or red fluorescence intensity was calculated from a boxed region surrounding the spine head. Spine volume was estimated using fluorescence from a cell fill (EGFP, tDimer-dsRed, or DsRedExpress)^70^. Bar graphs show the average values from the three time points occurring at 20-30 min after the most recent HFU (HFU_0_: 20-30min; HFU_30_: 50-60 min; HFU_45_: 65-75 min; HFU_60_: 80-90 min). Relative spine size was calculated by normalizing the individual spine fluorescence to the mean for all spines on the same dendrite using the two time points immediately prior to HFU.

### Statistics

All data are represented as mean ± standard error of the mean (SEM). All statistics were calculated across cells using GraphPad Prism. Statistical significance level (α) was set to p < 0.05 for all tests. When comparing only two groups, paired or unpaired (as appropriate) Student’s t-tests were used. When multiple comparisons were made, a one or two-way ANOVA with Bonferroni post-hoc test was performed. All p values are in the text and n values are in the figure legends.

## Acknowledgments

This work was supported by the National Institutes of Health (R01 NS062736, T32 GM099608, F99 NS125772) and an ARCS fellowship to JF. We thank J. Jahncke and L. Tom for support with experiments, J. Hell, J. Gray, J. Zheng, M. Navedo, S. Petshow, M. Anisimova, D. Sarkar, and M. Alarcon for critical input and/or reading of the manuscript.

## Figure Legends

**Figure S1.**
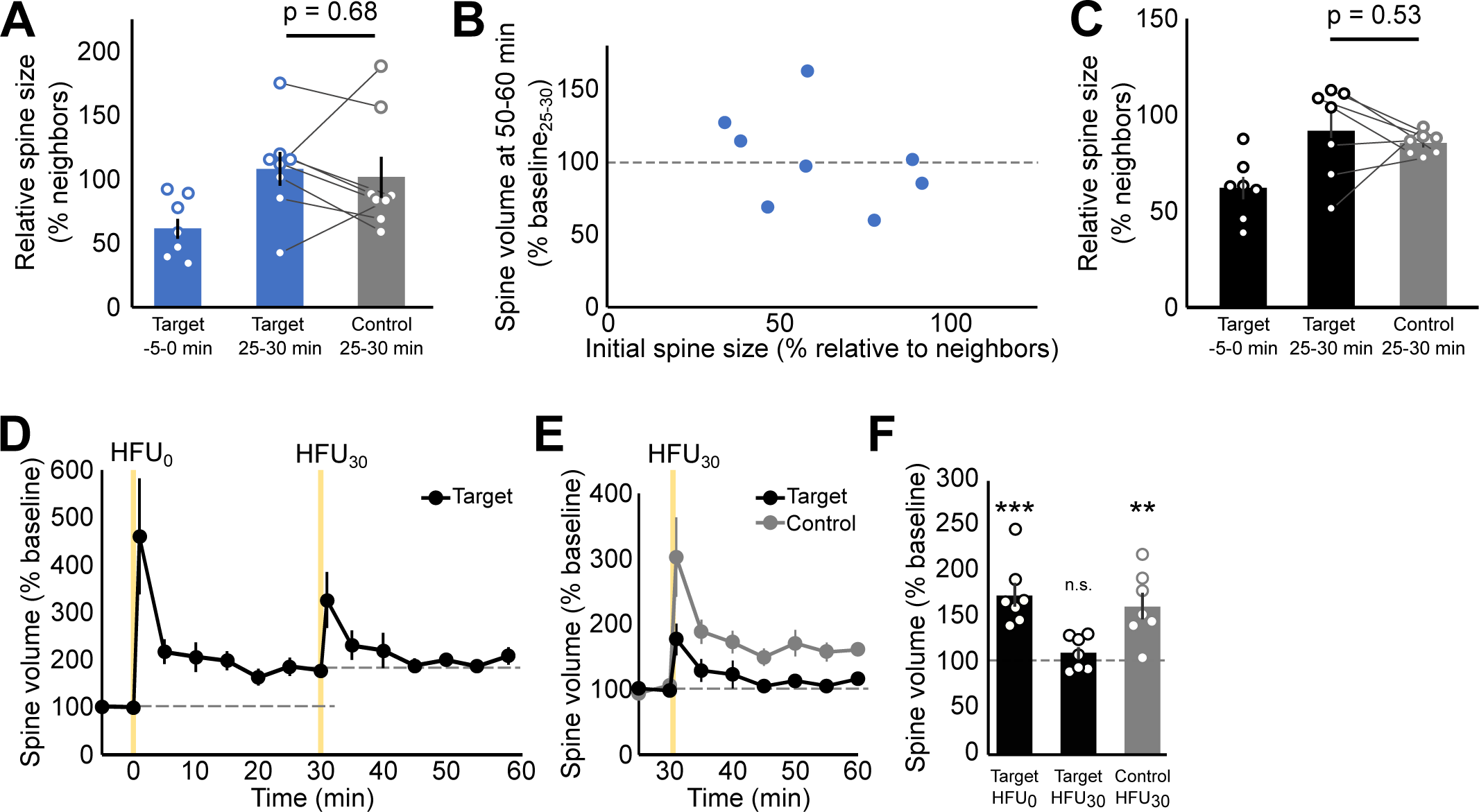
Related to Figure 1. **(A)** Quantification of spine size relative to neighbors for the target spine at baseline and 25-30 and control spine prior to HFU. There are no significant differences between the target’s post-stimulus and the control spine’s relative size (target: 108 ± 13%, control: 102 ± 16% ; p = 0.68). **(B)** Initial spine size did not correlate with saturation (p = 0.38). **(C)** The relative size of the control spine in the SEP-GluA2 and tD-dsRed condition is not significantly different to the target spine post-HFU (target: 92 ± 6%, control: 85 ± 2%,p= 0.53). **(D-F)** Spines from SEP-GluA2 experiments grew in response to HFU_0_ (measured as tD-dsRed fluorescence; black circles/bars; n = 7 spines/cells; 175 ± 13%; p<0.0001) but not HFU_30_ (111 ± 7%; p>0.99). Control spine grew in response to HFU_30_ (gray circles/bar; n = 7 spines/cells; 163 ± 14%). Statistics: paired Student’s t-test used in **A** and **C**; simple linear regression used in **B**. 2-way ANOVA with Bonferroni Test used in **F**.

**Figure S2.**
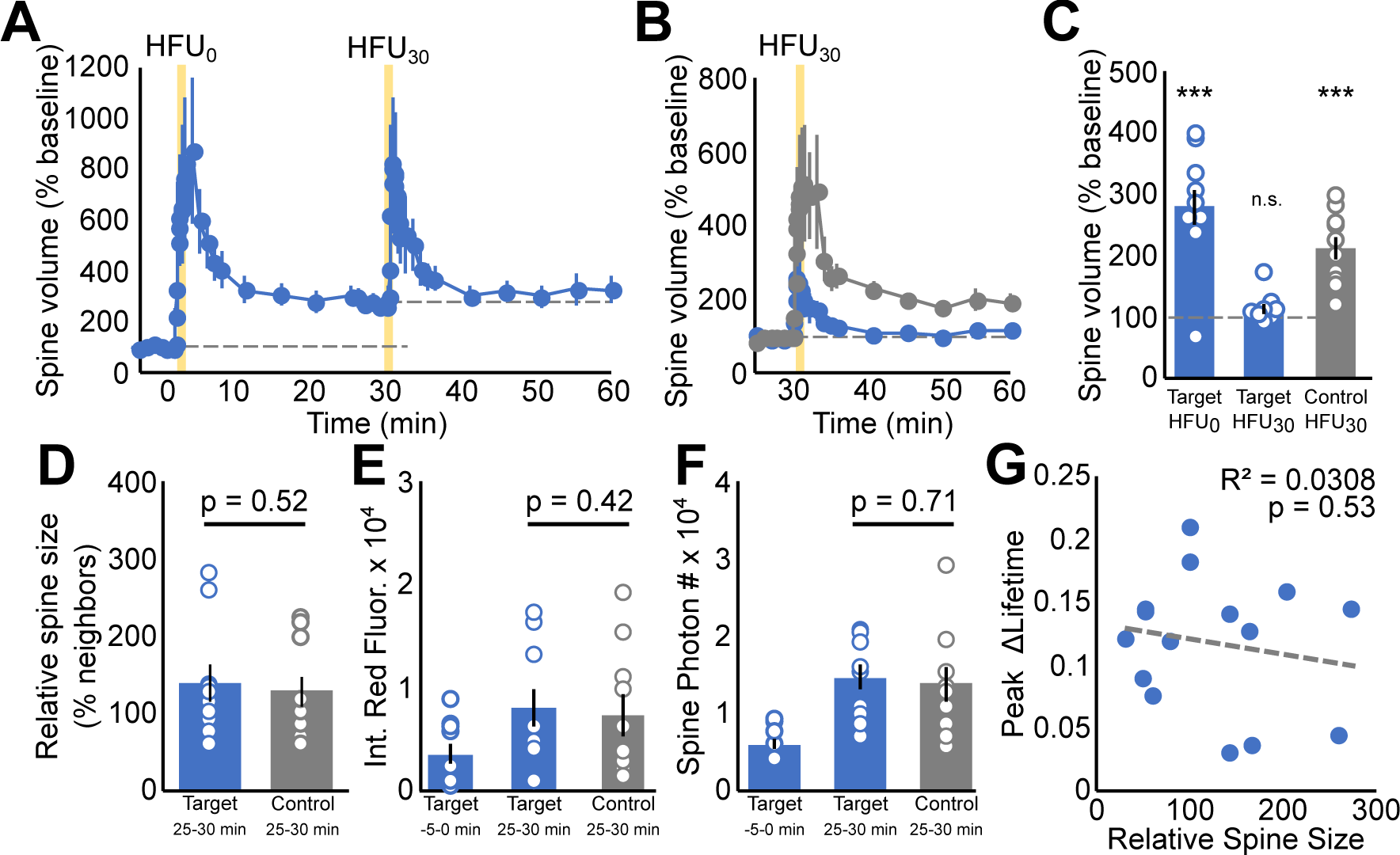
Related to Figure 2. **(A-C)** Target spines grew in response to HFU**_0_** (278 ± 30%; p < 0.001). The control spine but not target spines grew in response to HFU**_30_** (target: 113 ± 7%; p >0.99; control: 213 ± 19%; p < 0.001). **(D)** There was no significant difference in the relative size of the target after HFU**_0_** and the control spines (target: 139 ± 23%, control: 129 ± 19%; p = 0.52). **(E)** The integrated red fluorescence of the target after HFU and the control spine was not different (target: 78800 ± 17800, control: 72600 ± 20000; p = 0.42). **(F)** The number of photons collected in the target spine after HFU and the control spine was not different (target 14600 ± 1660, control: 13700 ± 2160; p = 0.71). **(G)** There was no correlation between spine size and CaMKII activation (p = 0.53). Data represent mean ± SEM. Statistics: 2-way ANOVA with Bonferroni test used in **C**; paired students t-tests used in **D-F**; linear regression analysis used in **G**. *p <0.05, 0.01 < **p < 0.001, ***p < 0.001.

**Figure S3.**
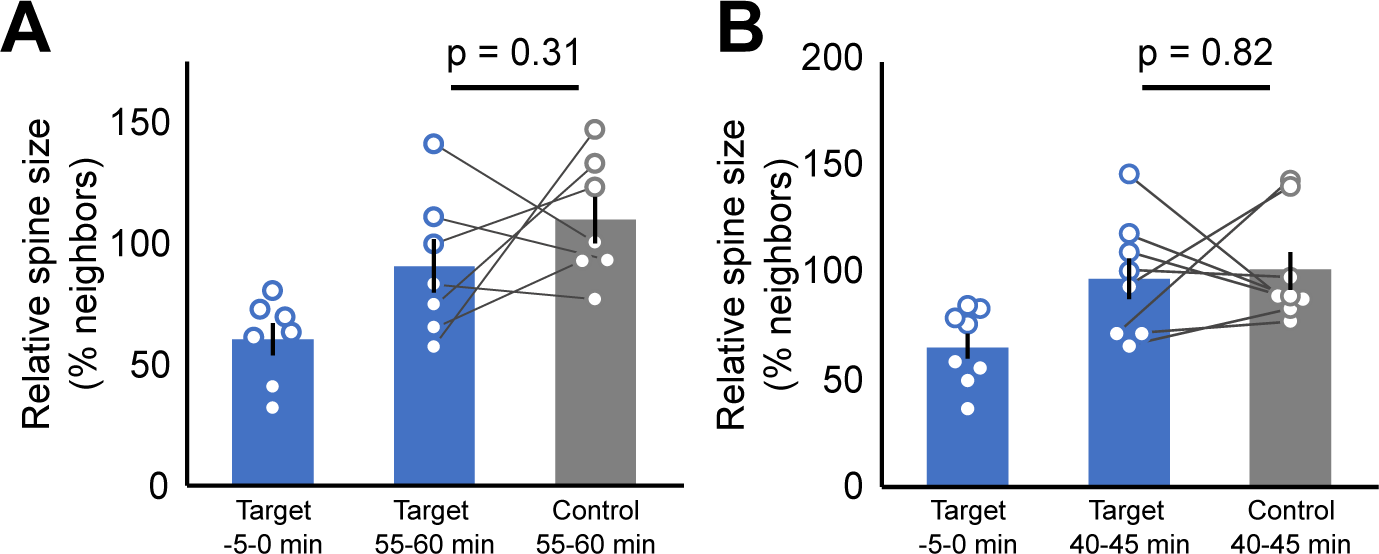
Related to Figure 3. **(A)** Quantification of spine size relative to neighbors for the target spine at baseline and 55-60 and control spine prior to HFU_60_. There are no significant differences between the target’s post-stimulus and the control spine’s relative size (target: 91 ± 11%, control: 110 ± 10%; p = 0.31). **(B)** Quantification of spine size relative to neighbors for the target spine at baseline and 40-45 and control spine prior to HFU_45_. There are no significant differences between the target’s post-stimulus and the control spine’s relative size (target: 96 ± 10%, control: 100 ± 9%; p = 0.82). Statistics: paired Student’s t-test used in **A** and **B.**

**Figure S4.**
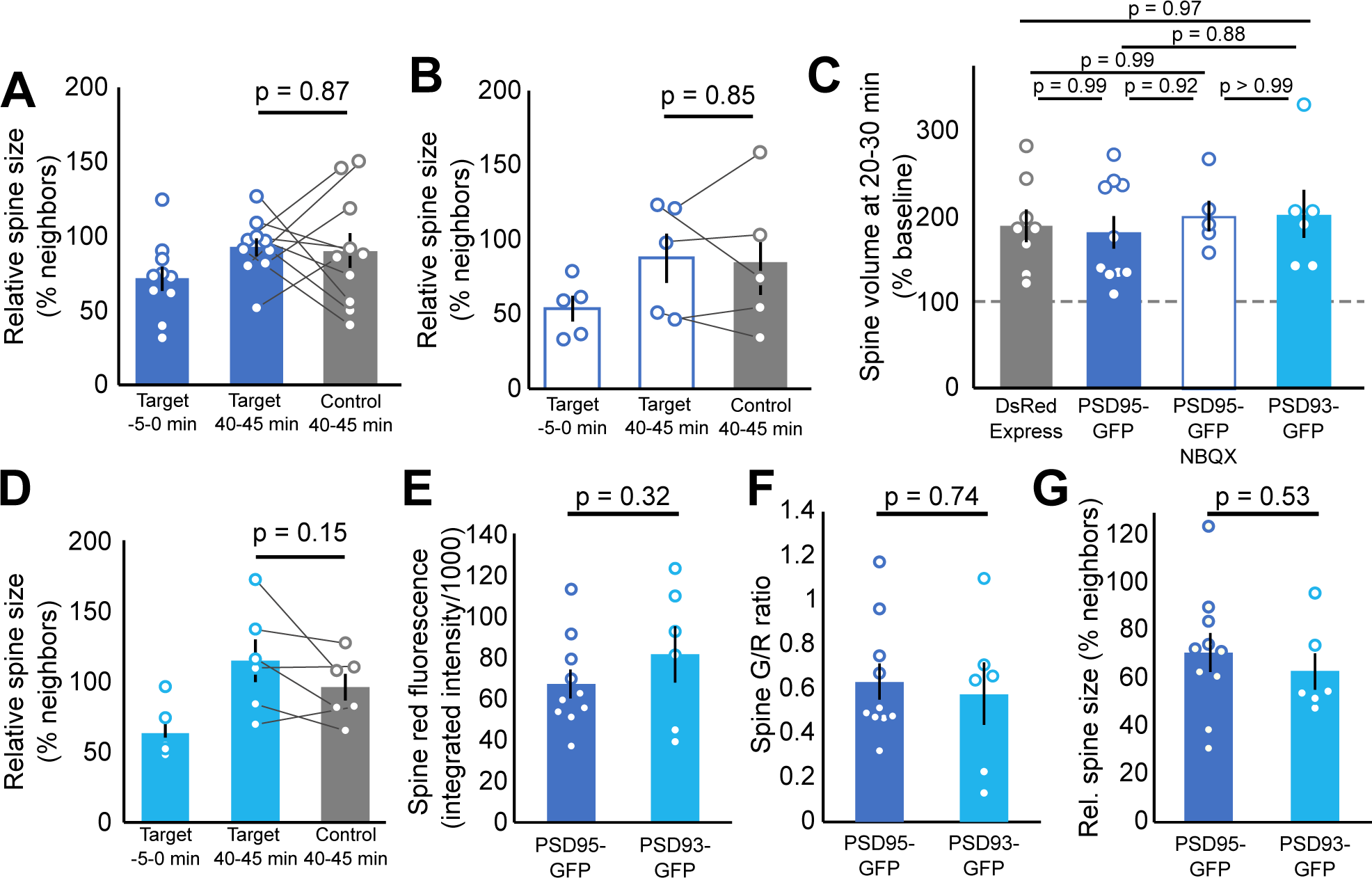
Related to Figure 4. **(A)** The post-HFU relative size of target spines from PSD95-GFP expressing cells was not significantly different from control spines (target 40-45 min: 93 ± 6%, control 40-45 min: 90 ± 12%, p = 0.87). **(B)** The post-HFU relative size of target spines from PSD95-GFP expressing cells treated with NBQX was not significantly different from control spines (target 40-45 min: 101 ± 10%, control 40-45 min: 96 ± 9%, p = 0.85). **(C)** The growth of the target spines in response to HFU_0_ was not significantly different from cells expressing only DsRedExpress (data from Figure 3) for any of the experimental groups (DsRedExpress: 189 ± 19%; PSD95-GFP: 180 ± 19%; PSD95-GFP + NBQX: 200 ± 18%; PSD93: 202 ±28; see bar graph for p-values). **(D)** The post-HFU relative size of target spines from PSD93-GFP expressing cells was not significantly different from control spines (target 40-45 min: 115 ± 15%, control 40-45 min: 96 ± 9%, p = 0.15). **(E)** The initial red fluorescence of the selected target spines was no different between the PSD95-GFP (67 ± 7) and PSD93-GFP (82 ± 14) conditions (p = 0.32). **(F)** The initial ratio of greent to red fluorescence of the selected target spines was no different (PSD95-GFP: 0.63 ± 0.08, PSD93-GFP: 0.58 ± 0.14, p = 0.74). **(G)** The initial relative size of target spines was no different between the groups (PSD95-GFP: 65 ± 5%, PSD93-GFP: 63 ± 8%, p = 0.53). Statistics: paired Student’s t-test used in **A, B, D**; ordinary one-way ANOVA with Bonferroni test used in **C**; unpaired Student’s t-test used in **E-F**.

